# Local rediscovery of the eucerine bee *Xenoglossa cressoniana* in Texas and Oklahoma after over a century of absence from the southern Great Plains

**DOI:** 10.64898/2026.05.22.727325

**Authors:** Elinor M. Lichtenberg, Isabelle S. Gonzales, Keng-Lou James Hung, John L. Neff

**Affiliations:** Department of Biological Sciences and Advanced Environmental Research Institute, University of Texas, Denton, TX, USA; Oklahoma Biological Survey, University of Oklahoma, Norman, OK, USA; Department of Chemistry and Biology, Toronto Metropolitan University, Toronto, ON, Canada; Central Texas Melittological Institute, Austin, TX, USA

**Keywords:** *Tetraloniella cressoniana*, Eucerini, longhorn bee, Great Plains, under-sampled region, *Salvia*

## Abstract

*Xenoglossa cressoniana*, also known as *Tetraloniella* cressoniana or *Xenoglossodes cressoniana*, is a eucerine bee known mainly from the US Great Plains. The species was described from a female collected somewhere in Texas in the early 1900s. Here, we report local rediscovery of this species in Texas and a new state record for Oklahoma after over a century with no intervening observations in either US state. While surveying north Texas ranches, we collected six specimens, including both males and females, at five sites northwest of the Dallas-Fort Worth Metroplex. In Oklahoma, we collected two females within a single nature preserve. *Xenoglossa cressoniana*’s range, the Great Plains and parts of the deep South, covers a large proportion of the United States that has been historically overlooked by most bee researchers. This report highlights the critical need for increasing data from under-sampled regions including the Great Plains and from private lands, even within a heavily sampled country such as the US.

## INTRODUCTION

Interest in pollinator declines and conservation has grown recently, yet the dynamics and biology of most individual species remain poorly studied. We report the local rediscovery of a rare bee with a broad geographic range. As described below, we focus on the region this species has historically been most associated with: the US Great Plains.

### Natural history

*Xenoglossa cressoniana* (Cockerell, 1905) is a solitary bee in the tribe Eucerini: the longhorn bees (Fig. 1). Like all other eucerines, *X. cressoniana* presumably excavates its nests in the ground (Danforth et al. 2019). Little is known about its natural history. It is probably oligolectic, collecting pollen mainly from *Salvia* species (Lamiaceae) including *S. azurea* Michx. ex Vahl and *S. reflexa* Hornem. (LaBerge 2001). It has also been found visiting *Boltonia, Helianthus microcephalus* Torr. & Gray, and *Helianthus petiolaris* Nutt. (all Asteraceae); *Peritoma serrulata* (Pursh) DC (Cleomaceae); *Phacelia* (Hydrophyllaceae); and *Verbena* (Verbenaceae) (LaBerge 2001, GBIF.org 2026). This species appears to be active in late summer and fall. Three specimens collected in Kansas and Idaho in the 1950s and ‘60s are reported as being collected in May (GBIF.org 2026). These could potentially be data entry or data management errors. They could also be specimen identification errors, although the Idaho specimen was identified by bee taxonomist Robert Brooks.

**Figure 1:**
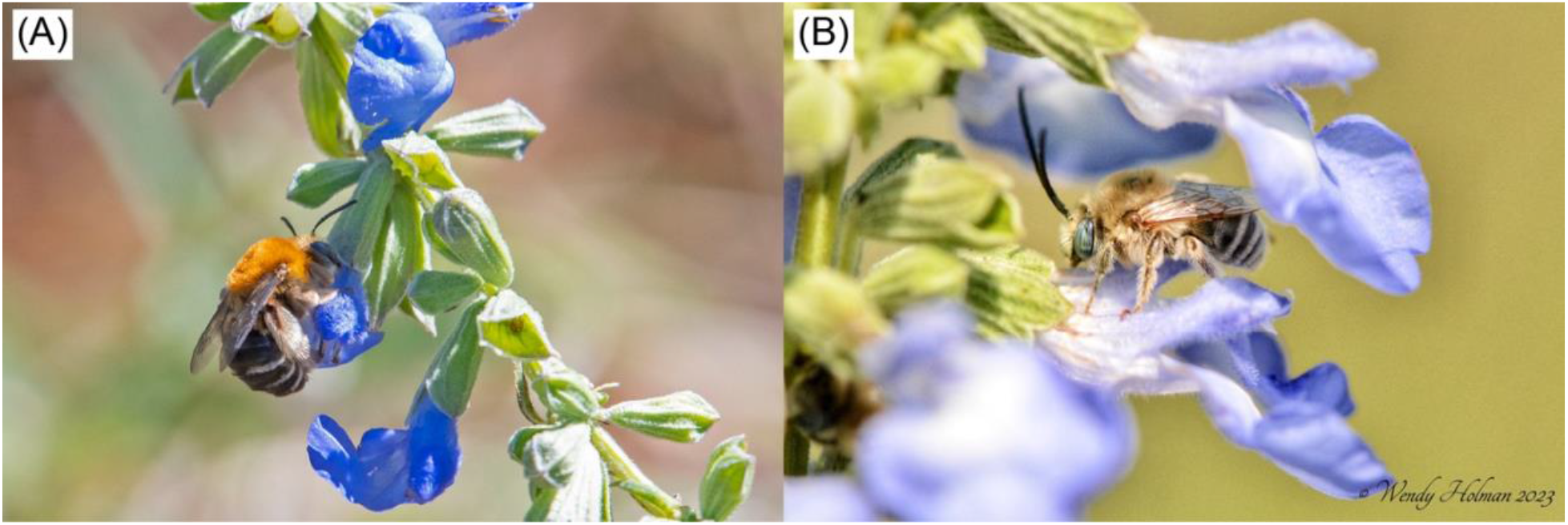
(A) Female and (B) male *Xenoglossa cressoniana* visiting sage flowers. Photos © (A) Lillie and (B) Wendy Holman, used under CC BY-NC license.

### Systematics

*Xenoglossa cressoniana* was first described as *Synhalonia cressoniana* based on a single female specimen collected somewhere in Texas, USA without any other information on the collector, date, or locale (Cockerell 1905). Like many other eucerine bees, *X. cressoniana* has had a complex taxonomic history. With the discovery of its male, *X. cressoniana* was transferred to *Tetraloniella* by LaBerge (2001) in a revision of what were considered to be the Western Hemisphere representatives of *Tetraloniella*. An alternate name used for the Western Hemisphere species placed in *Tetraloniella* is *Xenoglossodes* (LaBerge 2001). Recent molecular studies have reshuffled that picture. *Tetraloniella* was found to be a strictly Eastern Hemisphere group (Dorchin et al. 2018) and the Western Hemisphere species previously placed there were found to be a paraphyletic assemblage within an expanded concept of the genus *Xenoglossa* (Freitas et al. 2023). As the Western Hemisphere species previously placed in *Tetraloniella* (=*Xenoglossodes*) are highly paraphyletic and as *X. cressoniana* has not been included in any recent ph*y*logenetic study, the current position of *X. cressoniana* is uncertain (*incertae sedis*) within *Xenoglossa*.

### Geography

The type female of *Xenoglossa cressoniana* was collected in Texas. Later, Timberlake (1969) reported specimens from Nebraska and LaBerge (2001) from Alabama, Arizona, Colorado, Kansas, and New Mexico (Fig. 2 magenta circles; Table S1). These include the Temperate Grasslands, Savannas and Shrublands; and Temperate Conifer Forests biomes (Table 1).

**Table 1.** Biomes in which *X. cressoniana* has been reported, separated by country and state. Biomes follow The Nature Conservancy Terrestrial Ecosystems (The Nature Conservancy et al. 2009).

| **State/Province, Country** | **Biome(s)** | **Year Last Recorded** |
| --- | --- | --- |
| Alberta, Canada | Temperate Grasslands, Savannas and Shrublands | 2018 |
| Alabama, USA | Temperate Conifer Forests | Unspecified in (LaBerge 2001) |
| Arizona, USA | Temperate Conifer Forests | 1968 or Unspecified in (LaBerge 2001) |
| Colorado, USA | Temperate Conifer Forests | Unspecified in (LaBerge 2001) |
| Idaho, USA | Deserts and Xeric Shrublands | 1966 |
| Kansas, USA | Temperate Grasslands, Savannas and Shrublands | 2024 |
| Mississippi, USA | Temperate Conifer Forests | 2024 |
| Missouri, USA | Temperate Grasslands, Savannas and Shrublands | 2023 |
| Nebraska, USA | Temperate Grasslands, Savannas and Shrublands | 2025 |
| New Mexico, USA | Temperate Conifer Forests | Unspecified |
| Oklahoma, USA | Temperate Grasslands, Savannas and Shrublands | This study |
| Texas, USA | Temperate Grasslands, Savannas and Shrublands | This study |
| Hidalgo, Mexico | Deserts and Xeric Shrublands | 1962 |
| Michoacan, Mexico | Tropical and Subtropical Dry Broadleaf Forests | 1987 |

**Figure 2:**
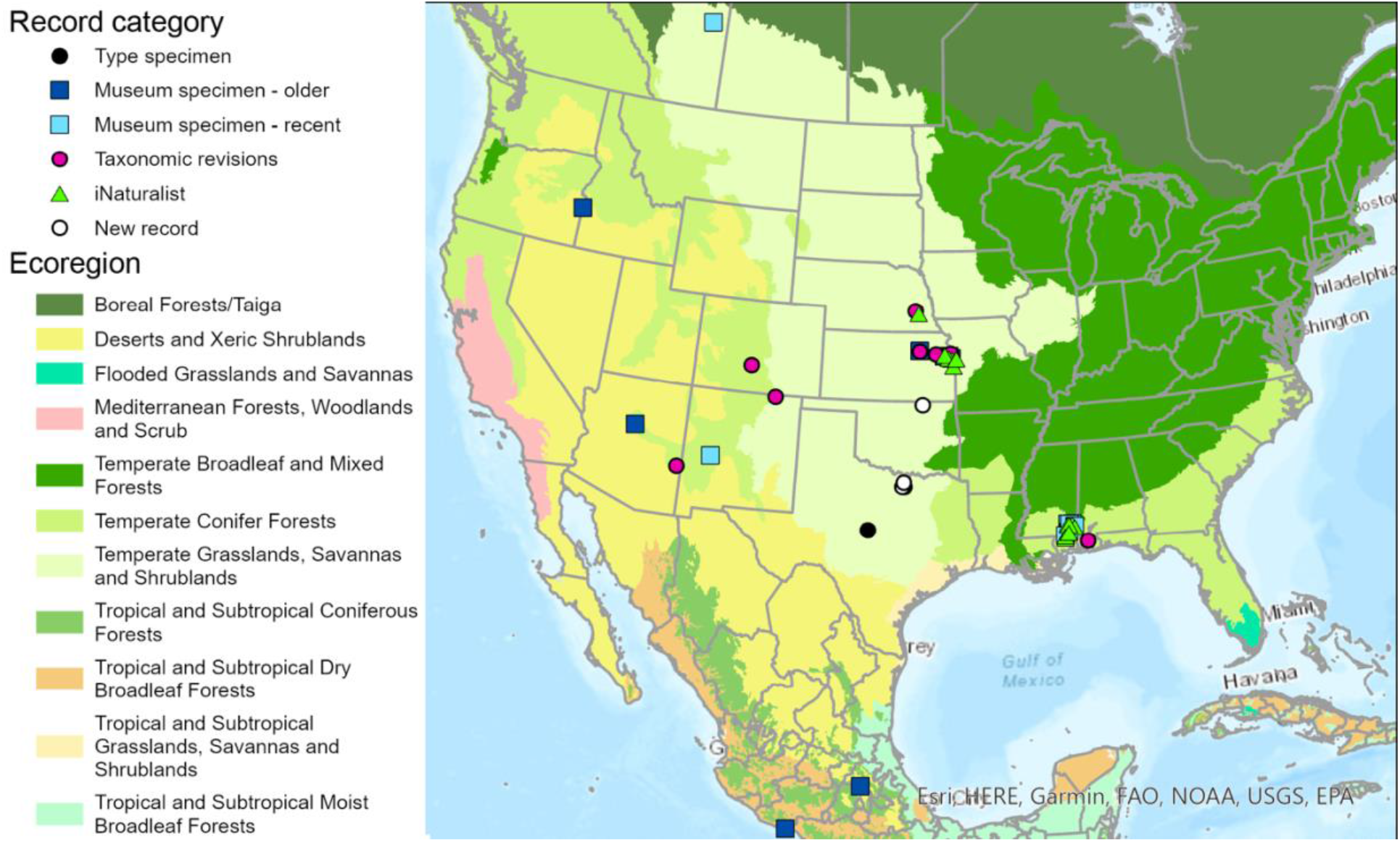
Map of known *X. cressoniana* records, assembled in ArcGIS Pro. Symbols indicate the record source, with museum specimens divided into older (1948-1982) and newer (since 2018). All iNaturalist observations were between 2018 and 2025. The type specimen’s collection location is reported only as Texas; we map it at the state’s geographic center. Taxonomic revisions are Timberlake (1969) and LaBerge (2001). Ecoregions are from The Nature Conservancy Terrestrial Ecosystems data layer (Olson and Dinerstein 2002).

We downloaded all *X. cressoniana* records currently available in GBIF as of February 2026, which are listed under *Tetraloniella cressoniana* (Fig. 2; Table S1). Twenty-one older records (before 1990) located this species in Arizona, Idaho, Kansas, Texas in the USA; and the Mexican states of Hidalgo and Michoacan. This includes the Deserts and Xeric Shrublands; Temperate Conifer Forests; Temperate Grasslands, Savannas and Shrublands; and Tropical and Subtropical Dry Broadleaf Forests biomes (Table 1). Fifteen museum specimens collected since 2018 were found in Alberta (Canada), Mississippi, and New Mexico. They are from the Temperate Conifer Forests; and Temperate Grasslands, Savannas and Shrublands biomes. iNaturalist contains 29 Research Grade observations from Kansas, Mississippi, Missouri, and Nebraska in the Temperate Conifer Forests; and Temperate Grasslands, Savannas and Shrublands biomes.

These records indicate the known range of *X. cressoniana* but also raise questions. KLJH examined the iNaturalist records and concurs with their identification. Few other fall-active Eucerini resemble our focal species due to the combination of all-black antennae, contrasting white apical and black basal or medial tergal hair bands, and the fulvous or ochraceous scutal pubescence of both sexes of *X. cressoniana*. In general, iNaturalist observations should be used with caution (Dickinson et al. 2010, Lowe et al. 2026) but do provide important clues to species’ distributions in regions that are under-studied or where most land is privately owned (Callaghan et al. 2021). This project did not have the scope to confirm the identification of all museum specimens. However, the *X. cressoniana* iNaturalist sightings and museum specimens report generally consistent locations. Several of the museum records aggregated in GBIF are disjunct from (Mexico, Idaho, Alberta) or are in distinctly different habitat (desert) from *X. cressoniana*’s previously reported range, which raises questions about this species’ range and biomes. Disjunct museum specimens collected in Mexico were identified by LaBerge in 2002, one year after he published a revised description of this species. The Alberta specimen was identified by bee taxonomist Lincoln Best using the dichotomous key in Laberge’s revision (2001), although this specimen diverged in several respects from individuals from *X. cressoniana*’s core range (L. Best, pers. comm.). The Idaho specimen and one Arizona specimen were identified by bee taxonomist Robert Brooks. This suggests that *X. cressoniana* as identified with current keys does have a fairly wide range that includes multiple biomes. However, the disjunct specimens should be critically re-evaluated, perhaps with genetic tools that can help refine species delineations (as in Rightmyer et al. 2014), before quantifying *X. cressoniana*’s range.

## STATUS

### Evidence

Here we report the local rediscovery of *X. cressoniana* in Texas and the first known records of the species from Oklahoma. In Texas, we collected two females and four males across five spatially independent sites that were separated by at least 1.5 km (Fig. 2 white circles; Table S1). These sites are located in Cooke, Denton, and Wise Counties in north Texas. In Oklahoma, we collected two females at a single patch of *Salvia azurea* at The Nature Conservancy’s Joseph H. Williams Tallgrass Prairie Preserve in Osage County, northeastern Oklahoma. Both states are in the southern Great Plains, which includes flatlands east of approximately the 96^th^ meridian in Texas, Oklahoma, and New Mexico. Texas specimens were collected in late September and early October of 2023 and Oklahoma specimens in mid-September of 2024. Specimens were identified by JNL (TX) and KLJH (OK) using LaBerge’s key and description (2001). *Xenoglossa cressoniana* appears to be widespread but patchy and uncommon, at present and perhaps even historically. Given how few specimens have been collected, and questions about the identity of some of those specimens, a formal assessment of extinction risk based on our local rediscovery is unlikely to be reliable at present.

### Search effort

We found the Texas *X. cressoniana* specimens while conducting research on private land in the Cross Timbers ecoregion, which is characterized by tallgrass prairies interspersed with dense woodland belts. In fall 2023, we sampled 19 sites during the fall bloom period which runs from late September into October. Additional sampling during the spring (April) and summer (June) bloom periods did not detect any *X. cressoniana*. Sites were “native pasture” ranches with largely native vegetation. At each site, we sampled pollinators through passive trapping and active netting on days with favorable weather: fully or mostly sunny, temperatures above 18° C, and average wind speed less than 6.7 m/s with gusts less than 8.9 m/s (as in Buckles and Harmon-Threatt 2019, Stein et al. 2020). We placed one blue vane trap at the site’s center and four sets of three pan traps 5 m from the blue vane trap for 24 h. Pan traps were 59 mL plastic bowls painted in colors that attract bees: Rust-oleum Fluorescent yellow and white, and Krylon bold neon blue (Kearns and Inouye 1993, Leong and Thorp 1999). Traps were placed at vegetation level, to reduce the known issue of traps attracting bees from long distances (Portman et al. 2020), and filled with soapy water. We additionally netted bees on flowers along eight 90 m transects radiating in cardinal and intercardinal directions from the site’s center. Netting occurred during two 45-min sessions that began at 10:00 and 13:00. All six specimens were found in blue vane traps with clear collection jars (BanfieldBio Inc.). Specimens are stored in the Lichtenberg Lab and will ultimately be deposited in the University of North Texas Elm Fork Natural Heritage Museum.

We found the Oklahoma *X. cressoniana* specimens while surveying bees, with a specific focus on bumble bees, at The Nature Conservancy’s Joseph H. Williams Tallgrass Prairie Preserve, a privately managed preserve open to the public and located within the Flint Hills ecoregion. Surveys at this preserve were part of a larger effort to implement the first-ever systematic state-wide survey of wild bees for Oklahoma. Standardized sampling was conducted across 96 sites in 2023 and 2024, with surveys at each site occurring during the late spring (May-June), summer (July-August), and early fall (September-October) of the same year for three total sampling events at each site. This timeframe primarily targeted bumble bee foraging activity. Study sites were located across all twelve level 3 ecoregions represented in Oklahoma, and varied in management practices. Each site consisted of a 50 m radius circle or, if located along a roadside, two parallel 100 m transects on opposite sides of the road. Sampling at each site occurred between 7:00 and 18:00 on days without rain. Specimen collection consisted of passive trapping using fluorescent blue vane and colored pan traps similar to those used in Texas, as well as targeted netting. We placed a blue vane trap at the center of the site and two sets of three pan traps immediately outside the survey area; traps were deployed for the duration of time we spent actively netting and collecting site-level habitat data. Surveyors performed 30 person-minutes of active bee netting by meandering across the entire survey area and examining patches of entomophilous blooms, prioritizing bumble bees whenever one was in sight. If plant species known to host pollen-specialist bees were blooming and abundant in the survey area, we performed supplemental netting on those blooms for an additional 5-15 min. Both specimens of *X. cressoniana* were captured on *Salvia azurea* during supplemental netting. One specimen has been deposited at the Sam Noble Oklahoma Museum of Natural History (SNOMNH); the other is temporarily stored in the Oklahoma Biological Survey’s bee reference collection but will ultimately also be deposited at SNOMNH.

### Threats

This species likely faces threats similar to other bees including habitat loss and simplification, parasites and pathogens, invasive species, pesticide exposure, climate change, and pollution (Potts et al. 2010, Goulson et al. 2015, Sánchez-Bayo and Wyckhuys 2019). *Xenoglossa cressoniana* appears to be oligolectic and thus threats to its primary host – bee-pollinated *Salvia* species (LaBerge 2001) – are of particular concern. Throughout *X. cressoniana*’s likely range, conversion of prairie habitat to developed areas and row crop agriculture, and increased frequency of extreme weather events, are likely particular threats (Heintzman and McIntyre 2019, Geng et al. 2023, Mehla and Jagadish 2025). Grassland habitat loss estimates within this species’ range are as high as 98% (Samson et al. 2004). Our Texas study area, at the southernmost end of the Great Plains, is experiencing significant conversion of natural areas to anthropogenic development (Liao et al. 2025). This area is on the outskirts of the Dallas-Fort Worth Metroplex, which is one of the largest and fastest-growing urban areas in the US (US Census Bureau 2025). Wise and Denton Counties are ranked 27^th^ (22% increase) and 60^th^ (18%), respectively, in fastest growth of all US counties between 2020 and 2025 (US Census Bureau 2026).

The fact that we found multiple *X. cressoniana* specimens with limited sampling effort in Texas highlights an additional threat to both science and conservation planning: significant under-sampling of the Great Plains and private lands. Researchers studying a range of taxa have noted that the Great Plains receive significantly less attention than other biomes (e.g., odonates Reece and McIntyre 2009, spiders Nemec 2014, bees Jamieson et al. 2019, insects in general Bulger et al. 2026) despite covering approximately one third of the contiguous United States. A more systematic investigation suggests that bee data completeness for most of the Great Plains is only 10-20% (despite a history of bee research in the central Great Plains), and calls for greater sampling and specimen digitization in the central and southern US (Chesshire et al. 2023). Private lands comprise the majority of land globally, including 96% of land in Texas (Drescher and Brenner 2018) and 97% of land in Oklahoma (York and Jager 2021). Private lands are also significantly under-sampled (Hilty and Merenlender 2003) due to the increased effort needed to gain access or obtain permission to collect specimens on private lands (Hargiss and DeKeyser 2014) even when they are open to the public as the Joseph H. Williams Tallgrass Prairie Preserve is. However, private lands can be important in conserving biodiversity (Nolasco et al. 2026).

Documenting such biodiversity both through community science (Callaghan et al. 2021, Gillard and Rowley 2026) and more formal sampling is necessary to achieve a more complete understanding of the current status of many uncommon species including *X. cressoniana*. Such under-sampling delays species discovery (e.g., Bossert et al. 2024, 2026), leads to incorrect biodiversity estimates and range maps (e.g., Gibson et al. 2018, Carper et al. 2019), and limits ability to capture range expansions or contractions (Martens et al. 2022). These inaccuracies, in turn, limit predicting impacts of specific threats and understanding which species are most at risk (reviewed in Jamieson et al. 2019).

## CONCLUSION

This report highlights the critical need for more data from under-sampled regions including the Great Plains. We found six *X. cressoniana* specimens with limited sampling in north-central Texas, suggesting that this species may be less uncommon in that region. Due to lack of sampling, however, the species had remained unreported in Texas for over a century. In contrast, the species may be genuinely rare in Oklahoma, as we found *X. cressoniana* only once there despite standardized surveys at nearly 100 locations throughout the state. The detection location was very close to the border with Kansas, where a third of sightings of this species were reported. Expanding sampling efforts beyond bee researchers’ established sites, sampling more private lands, and increasing use and utility of community science data are critical for understanding species ranges, conservation status, most pressing threats, and potentially effective conservation actions.

## DATA ACCESSIBILITY

The GBIF search (GBIF.org 2026) and the Symbiota collection containing our new Texas records (Lichtenberg 2026) are publicly available. Our new Oklahoma records are in Table S1. Other data points come from published records (Timberlake 1969, LaBerge 2001).

## ACKNOWLEDGEMENTS

We thank V. Dietrich, I. Lugo, J. Martinez, J. Moore, M. Muñiz, S. O’Dell, A. Pearson, and E. Phillips for field sampling and specimen processing; J. Ascher for his contributions to identifying bees on iNaturalist; and two anonymous reviewers and the journal editors for feedback. This material is based upon work that is supported by the National Institute of Food and Agriculture, U.S. Department of Agriculture, under award number 2022-38640-37488 through the Southern Sustainable Agriculture Research and Education program under subaward number OS23-162; and the Oklahoma Department of Wildlife Conservation, under Federal Aid Grant number F22AF03005 through the State Wildlife Grants program, award number OK T-131-R-1. USDA is an equal opportunity employer and service provider.

